# Are genetically defined “metapopulations” self-evident in YHRD?

**DOI:** 10.64898/2026.02.07.704579

**Authors:** Tóra Oluffa Stenberg Olsen, Mikkel Meyer Andersen, James Curran, Michael Krawczak, Amke Caliebe

## Abstract

In forensic genetics, the evidential value of a match between the Y-chromosomal short tandem repeat (Y-STR) profiles of a trace and a suspect is typically quantified by the frequency of the profile in a population database, particularly the Y-chromosomal Haplotype Reference Database (YHRD). However, for this approach of obtaining a ‘match probability’ to be valid, the database population must be representative of all plausible alternative trace donors in a given case. Since appropriately defining such a ‘suspect population’ can be difficult, YHRD highlights so-called ‘metapopulations’ that comprise profiles from different, geographically dispersed populations with presumed shared ancestry. We investigated whether such metapopulations are self-evident in the current version of YHRD. To this end, we performed classical cluster analysis using allele dissimilarity as a measure of pairwise distance between Y-STR profiles. Our analyses revealed only a weak genetic structure in YHRD the extent of which was inversely proportional to the respective marker mutation rate. This suggests that YHRD cannot be divided into clearly distinguishable subgroups based solely on the genetic information it contains, at least not into subgroups that would correspond closely to the metapopulations highlighted in the database itself. If profile frequencies in metapopulations are to continue to be equated with match probabilities, then a clearer definition of metapopulations and a better justification of their use in forensics are needed.

## 1. Introduction

The analysis of Y-chromosomal short tandem repeat (Y-STR) markers plays an important role in forensic genetics, particularly in cases, such as sexual assaults, where the biological trace of interest contains an unbalanced mixture of a major female and a minor male DNA component. Due to their male specificity, single-source Y-STR profiles can often still be reliably determined from mixtures even when the ratio of male to female genetic material is minimal [1, 2].

Once a suspect has been identified in cases like the above, his Y-STR profile can be determined as well and compared to the trace profile. While a mismatch between the two profiles normally excludes the suspect from being a donor, matches are more difficult to interpret. This is because, in court, the forensic genetic expert must quantify the strength of evidence of a match, usually in the form of the probability with which such a match would also be observed between the trace profile and the profile of another male. The question of how best to obtain this so-called ‘match probability’ and, in particular, what “another” should mean in this context has occupied forensic geneticists for more than two decades [3–5].

The most commonly used approach to determining a match probability equates the latter with the frequency of the Y-STR profile of interest in a suitable group of plausible alternative donors also referred to as the ‘suspect population’ [5, 6]. Although various mathematically sound methods have been proposed to estimate a haplotype frequency from a representative sample of a suspect population [7–9], the main concern to date has been how to adequately define this population itself. Notably, the DNA commission of the International Society for Forensic Genetics (ISFG) recommended use of the geographical location of the crime scene as a “leading criterion” for selecting the “right” suspect population [2].

The ISFG proposal is certainly meaningful because one of the key criteria for the plausibility of donorship is spatial proximity to the place where the trace was recovered. It is undoubtedly reasonable to imply that most plausible alternative donors lived near the crime scene at the time. However, with very few exceptions, estimating the population frequency of a Y-STR profile requires a sample or database that is both sufficiently large and sufficiently representative of the population in question. ‘Representative’ here takes a statistical definition and means that the sample or database adequately captures all relevant population characteristics. The largest database established for this purpose is the Y-chromosomal Haplotype Reference Database (YHRD), which emerged in 2004 from a series of previous efforts in Europe [10]. Since then, YHRD has been continuously extended and currently contains approximately 350,000 Y-STR profiles from all over the world, comprising between eight and 27 Y-STR markers.

Despite these tremendous efforts, even a database the size of YHRD cannot cover every case-relevant geographical target region at a level (village, city, state, province, etc.) desirable for forensic casework. Moreover, humans have undergone a multitude of demographic processes that led to the formation, and subsequent reorganization, of local populations at varying levels of intra- and inter-group genetic differentiation. To allow forensic experts to take this fact into account when using YHRD, the database curators decided to highlight “metapopulations” in YHRD. These are defined as combinations of originally separate, regional subsets of the data that were supposedly connected by past migration or gene flow [10]. The stated goal of the endeavour has been to group together “geographically dispersed human population samples with shared genetic ancestry” [2], using geographic, linguistic, and demographic bonds as proxies for (male) genetic relatedness.

In the present study, we aimed to assess the extent to which the above-mentioned objective was achieved. To this end, we investigated

i. whether clusters of Y-STR profiles are evident in YHRD that indicate subsets with markedly more recent coancestry,
ii. to what extent the degree of clustering observed for different marker combinations depends upon their mutation rates, and
iii. whether the Y-STR profile clusters thus identified overlap with the metapopulations defined in YHRD.

Ideally, our study would have been based upon an *ab initio* reconstruction of the genealogy of the Y-STR profiles included in YHRD, using software tools publicly available for this purpose. However, the reconstruction of coalescence trees is not only computationally intensive, but its results also critically depend upon the underlying evolutionary model and the choice of model parameters [11]. Therefore, we performed classical cluster analysis using marker-wise allelic dissimilarity as an (agnostic) measure of the pair-wise distance between Y-STR profiles. Although this approach may not perfectly reproduce the results of a coalescence-based analysis, it should still yield trees that adequately reflect the true genealogy of the Y-STR profiles included in YHRD.

## 2. Methods

### 2.1. Data and data analyses

All data analyses were carried out using R [12] version 4.5.0 and the packages `rforensicbatwing` [13] version 1.3.1, `umap` [14] version 0.2.10.0, `Rtsne` [15–17] version 0.17, `cluster` [18] version 2.1.8, `fields` [19] version 16.3.1, `LEA` [20] version 3.16.0, `tidyverse` [21] version 2.0.0, `pals` [22] version 1.10, `dendextend` [23] version 1.19.0,`fpc` [24] version 2.2.13, `ggpubr` [25] version 0.6.0, `seriation` [26, 27] version 1.5.8, `data.tree` [28]version 1.1.0, `igraph` [29, 30] version 2.1.4, `ggraph` [31] version 2.2.1, and `cowplot` [32] version 1.1.3. The respective code is publicly available online [33].

We used data from YHRD [34] in accordance with a project proposal previously submitted to the database curators (https://yhrd.org/pages/Projects/P5). Currently, YHRD defines seven major metapopulations, namely African, Afro-Asiatic, Native American, Australian Aboriginal, East Asian, Eskimo-Aleut, and Eurasian, in addition to one ‘Admixed’ metapopulation covering populations with pronounced admixture [2, 34]. Some of these YHRD metapopulations are further divided into subgroups (Fig. 1) that YHRD also refers to as ‘metapopulations’, but from which we distinguish the original metapopulations by adding the term ‘major’ to the latter, if required. A more detailed description of the geographical range of the YHRD metapopulations can be found in Table 2 in [2].

**Fig. 1:**
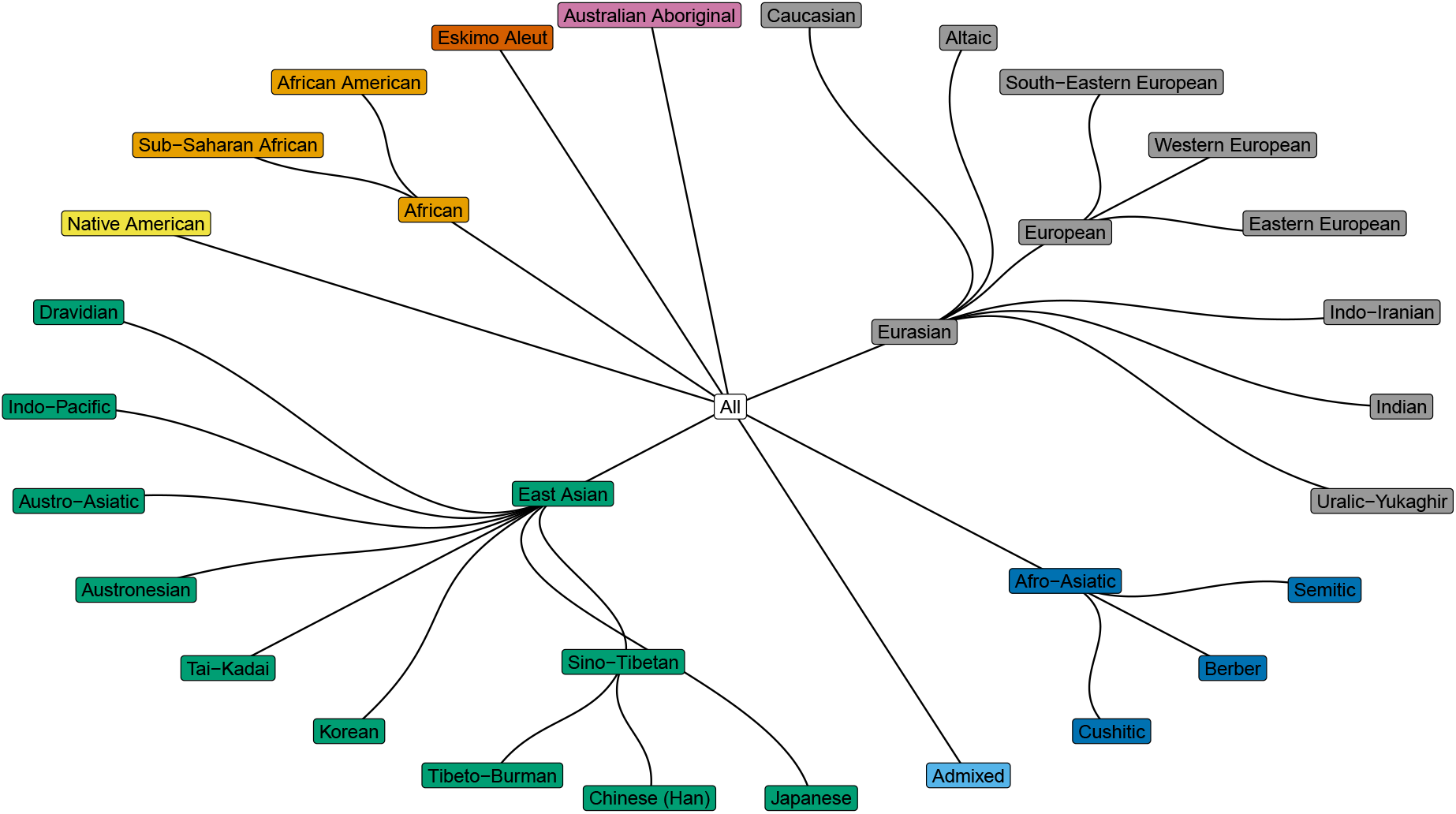
Overview of YHRD metapopulations. Colors correspond to the major YHRD metapopulations defined in [34], including the ‘Admixed’ metapopulation.

We generated four sets of Y-STR loci henceforth referred to as ‘kits’. Each kit consisted of eight Y-STR loci that were selected based upon their mutation rates (Table 1). We refer to these kits as ‘fast’, ‘medium’, ‘mixed’ and ‘slow’ depending on the mutation rates of the loci included in the respective kit. Only integer-valued alleles were considered here because distances to and between intermediate alleles are difficult to define in accordance with evolutionary proximity. The numbers of Y-STR profiles for each YHRD metapopulation and each kit are provided in Supplementary Table S.1.

**Table 1.** Mutation rate-dependent definition of Y-STR marker kits.

| Kit | Fast | Medium | Mixed | Slow |
| --- | --- | --- | --- | --- |
| **Mutation rate<sup>a</sup>** | $4.36 \times 10^{-3} - 1.40 \times 10^{-2}$ | $2.34 \times 10^{-3} - 4.34 \times 10^{-3}$ | $3.95 \times 10^{-4} - 1.40 \times 10^{-2}$ | $3.95 \times 10^{-4} - 2.24 \times 10^{-3}$ |
| **Loci** | DYS627, DYS576, DYS518, DYS449, DYS570, DYS458, DYS439, DYS460 | DYS481, DYS549, DYS456, DYS533, DYS635, DYS389I, YGATAH4, DYS391 | DYS627, DYS576, DYS518, DYS449, DYS438, DYS392, DYS448, DYS437 | DYS438, DYS392, DYS393, DYS437, DYS643, DYS448, DYS390, DYS19 |
| **Number of Y-STR profiles<sup>b</sup>** | 102,253 | 102,880 | 103,129 | 107,748 |
| **Number of YHRD metapopulations** | 25 | 27 | 25 | 27 |

<sup>a</sup>Mutation rates were taken from www.yhrd.org. <sup>b</sup>Numbers include only Y-STR profiles with integer-valued alleles.

**Table 2.** Agglomeration coefficients (*ac*) obtained with different agglomeration methods.

| Marker kit<sup>a</sup> | Agglomeration method |  |  |  |  |
| --- | --- | --- | --- | --- | --- |
|  | Average | Single | Complete | Ward | Weighted |
| **Fast** | 0.89 | 0.80 | 0.93 | 0.99 | 0.90 |
| **Medium** | 0.93 | 0.92 | 0.96 | 0.99 | 0.95 |
| **Mixed** | 0.91 | 0.84 | 0.95 | 0.99 | 0.93 |
| **Slow** | 0.97 | 0.97 | 0.98 | 1.00 | 0.98 |

<sup>a</sup>For the definition of marker kits, see main text and Table 1.

Except for the analysis explained in Section 2.3, the number of Y-STR profiles was limited to a maximum of 100 per YHRD metapopulation for each marker kit to ensure a balanced ancestry of the resulting sample. The Y-STR profiles were selected randomly from the YHRD metapopulation under consideration.

### 2.2. Spatial autocorrelation

We examined the degree of spatial autocorrelation between Y-STR profiles and their sampling locations, using Moran’s *I* [35]. In particular, we repeated the analysis by Roewer et al. [36] to assess whether similar levels of autocorrelation as observed in early YHRD also characterise the data analysed in the present study. First, we calculated pair-wise Great Circle Distances (in kilometres) between sampling locations from the respective longitude and latitude data. Y-STR profile pairs were then grouped into distance classes defined as adjacent 250 kilometres-wide intervals, with one additional class comprising all pairs sampled at the same location. We then calculated Moran’s *I* for each class, setting the required spatial weights for Y-STR profile pairs equal to unity, if the profile pair fell into that class, and equal to zero otherwise. While a value of *I* close to plus one indicates similarity of Y-STR profiles in a given distance class, and a value of *I* close to minus one indicates dissimilarity, *I* = 0 means a lack of correlation in repeat number between Y-STR profiles in the respective class [35].

### 2.3. Distance measures

The possible structuring of the YHRD data was to be analysed by multi-dimensional scaling (MDS) and hierarchical clustering, which requires a pair-wise distance measure between Y-STR profiles. For this, we considered the following three quantities:

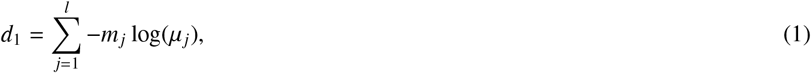

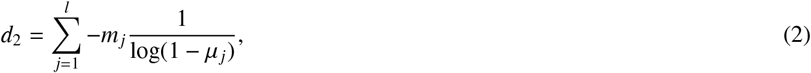

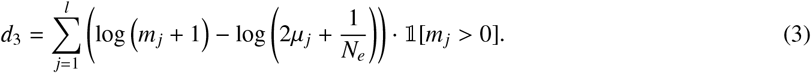

Here, *l* is the number of loci, *µ*_*j*_ denotes the mutation rate at the *j*^th^ locus, *m*_*j*_ is the absolute difference in repeat number, at the *j*^th^ locus, between the two Y-STR profiles of interest, and *N*_*e*_ denotes the effective size of the underlying population. We considered effective population sizes of 1,000, 5,000, and 10,000, respectively [37]. All three quantities satisfy the properties of distance measures (Appendix A).

The rationale of all three distance measures was to consider the respective mutation rates when weighting the allelic differences. Hence, we introduced locus-specific weights that depended differently upon each locus-specific mutation rate. More precisely, *d*_1_ weights the repeat difference by − log(*µ*_*j*_) so that loci with a low mutation rate contribute disproportionally to the overall profile dissimilarity. The second distance measure, *d*_2_, weights the repeat difference with −1*/* log(1 − *µ*_*j*_), which has a similar effect to *d*_1_ but gives even more weight to loci with lower mutation rates. The third measure, *d*_3_, additionally considers the estimated time to the most recent common ancestor of the two profiles of interest. Thus, *d*_3_ subtracts log(2*µ*_*j*_ +1*/N*_*e*_) from log(*m*_*j*_ +1) to simultaneously account for the divergence of profiles expected from mutation and drift. A more detailed justification of *d*_3_ is provided in Appendix B. In addition to *d*_1_ to *d*_3_, we also considered the Euclidean and Manhattan distance.

All distance measures used in the present study were computational compromises that likely resulted in some loss of ancestry information. Therefore, we compared each measure to BATWING [38] estimates of the total length of the coalescent branches between two Y-STR profiles, as described in [13]. Unfortunately, due to high computational costs, BATWING estimates were not suitable as distance measures themselves. For each marker kit, we instead selected 30 Y-STR profiles at random, estimated the lengths of the coalescent branches for all possible profile pairs, and repeated this procedure five times. The other distance measures were then related to the BATWING estimates in each repetition by way of Spearman’s rank correlation coefficient.

### 2.4. Dimensionality reduction

To be able to visualise whether and how Y-STR profiles from YHRD group into clusters, we first employed three methods of dimensionality reduction, namely MDS, Uniform Manifold Approximation and Projection (UMAP), and t-Distributed Stochastic Neighbour Embedding (t-SNE) [16, 17, 39–41]. The three methods differ in how they maintain the overall structure of distance in the data during dimensionality reduction. While MDS aims to preserve all pairwise distances between Y-STR profiles as much as possible, t-SNE focuses on the maintenance of local relationships. Therefore, t-SNE often produces well-separated clusters, but only inadequately captures their global relationships. UMAP lies somewhere in between and typically yields similar results to t-SNE regarding local structure, but tends to infer the global structure somewhat better than t-SNE, although still less reliably than MDS.

### 2.5. Clustering

For each marker kit, the corresponding Y-STR profiles were subjected to hierarchical clustering with one of five possible agglomeration methods: ‘average’, ‘single’, ‘complete’, ‘Ward’, and ‘weighted’ [42]. The selection of the agglomeration method was based upon the respective agglomeration coefficient, *ac*. This figure lies between zero and unity and relates the average dissimilarity of a Y-STR profile to the first cluster it joins to the dissimilarity between the last two merged clusters [18]. An *ac* value close to unity indicates a data structure where data points join clusters early, relative to the final merger, thereby suggesting the presence of well-defined and tight clusters. It should be noted that *ac* tends to increase with an increasing number of observations and therefore should not be compared between samples of different sizes.

For a given marker kit, we performed hierarchical clustering with the agglomeration method that yielded the largest *ac* and with the distance measure that correlated most strongly with the coalescent tree branch lengths estimated before with BATWING. The number of clusters present in the resulting tree was determined using the elbow method [43]. By plotting a given tree height against the number of sub-clusters present at that height, the elbow method identifies a point (the ‘elbow’) where the decrease in height per added cluster slows down significantly. This property suggests that adding more clusters beyond the elbow only slightly increases the similarity of Y-STR profiles within clusters.

The stability of the inferred clusters was assessed by bootstrapping using the Jaccard index [24, 44–46] as a measure of cluster overlap. For this purpose, 100 bootstrap samples were drawn ignoring multiple drawings of one and the same Y-STR profile. Each bootstrap sample was then subjected to the same hierarchical cluster analysis as the original data and decomposed into the same number of clusters as originally inferred for the respective marker kit. For each original cluster, the most similar bootstrap cluster was then identified through maximisation of the pairwise Jaccard index. Finally, these maximum Jaccard indices were averaged over all bootstrap runs to yield a measure of stability of each original cluster.

In addition to hierarchical clustering, we also performed STRUCTURE [47] analyses as a means to group Y-STR profiles. STRUCTURE does not aim to maximise Y-STR profile similarity within groups, but rather maximises the overall probability of each profile being assigned to a specific group by varying the profile frequencies within groups. STRUCTURE analyses were performed in R as described in [48, 49]. The marker kit-specific numbers of groups were again determined using the elbow method [43], but this time employing a cross-entropy criterion [49] calculated for between one and 15 groups present.

## 3. Results

### 3.1. Spatial autocorrelation

Spatial autocorrelation analysis with Moran’s *I* index revealed that the current relationship between Y-STR profile characteristics and sampling location resembled that reported for an early version of YHRD [36]. Thus, for each marker kit, Y-STR profiles tended to be more similar (i.e. Moran’s *I >* 0) when sampled at nearby rather than distant locations, and increasing the Great Circle Distance consistently decreased the spatial autocorrelation (Fig. 2).

**Fig. 2:**
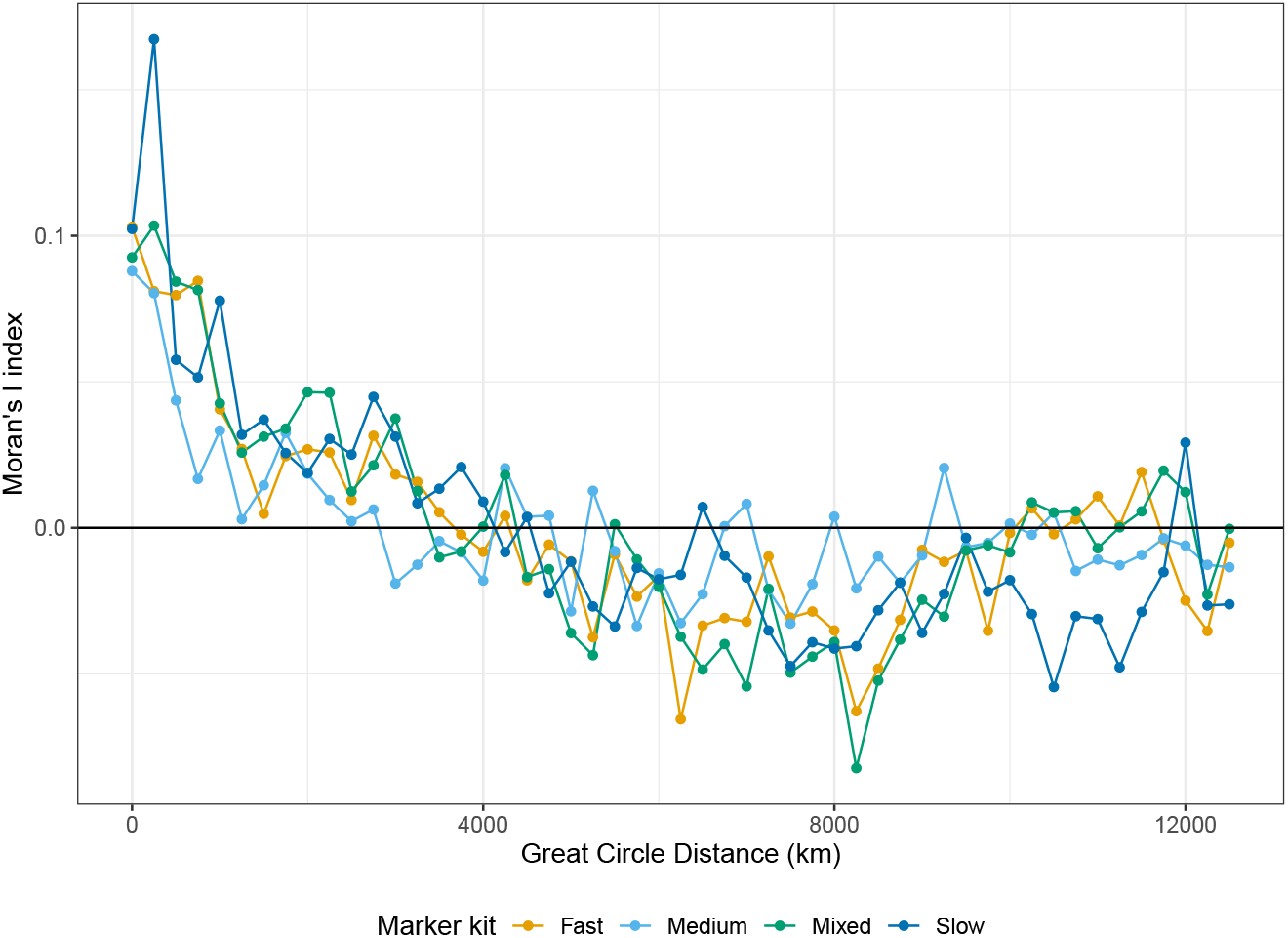
Spatial autocorrelation (Moran’s *I* index) analysis of Y-STR profiles. For the definition of marker kits, see main text and Table 1.

### 3.2. Distance measures

To select the most appropriate of five possible distance measures for further analysis (see Section 2.3), each distance measure was compared to the coalescent tree branch lengths estimated with BATWING [13, 38], assuming an effective population size of 1,000 (Fig. 3; for results obtained with effective population sizes of 5,000 and 10,000, see Supplementary Figs. S.1 and S.2, respectively). Benchmarking against BATWING was performed in five repetitions, separately for each marker kit.

**Fig. 3:**
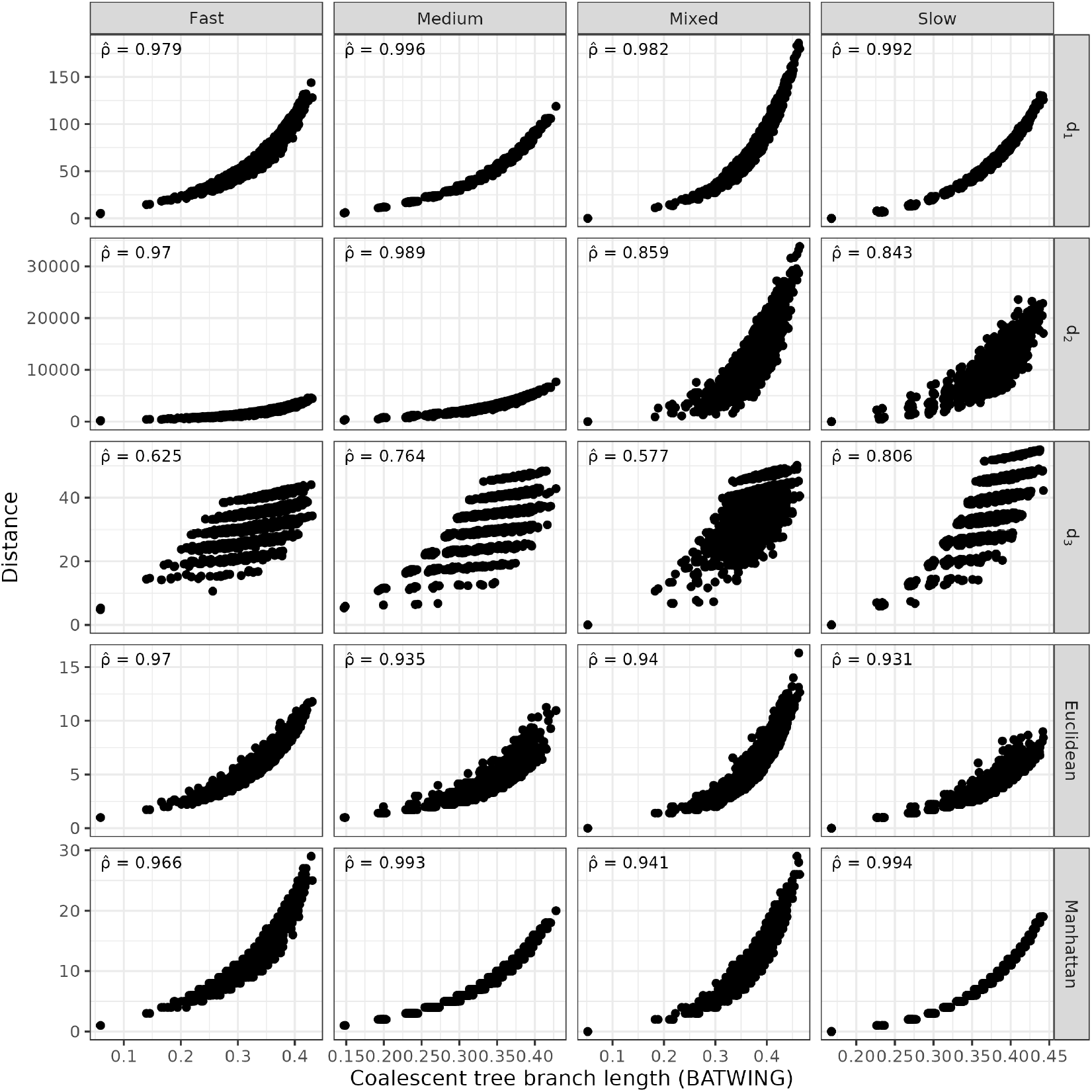
Benchmarking of pairwise Y-STR profile distance measures against BATWING. For each pair of Y-STR profiles, the respective distance is plotted against the coalescent branch length estimated with BATWING, assuming an effective population size of 1,000. 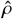: average Spearman’s rank correlation coefficient, taken over five repetitions of the analysis. For the definition of marker kits, see main text and Table 1.

We selected *d*_1_ as the most appropriate distance measure for further analysis for all marker kits because it consistently yielded the strongest correlation with the coalescent tree branch lengths (Fig. 3). Interestingly, the results obtained for *d*_1_ were very similar to those obtained for the Manhattan distance. This suggests that additional weighting of the absolute allele difference for a given Y-STR by minus the logarithm of the corresponding mutation rate (see Section 2.3) did not significantly affect distance measure performance for the marker kits considered.

### 3.3. Dimensionality reduction

Employing distance measure *d*_1_, the Y-STR profiles were next subjected to MDS analysis separately for each marker kit (Fig. 4). If Y-STR profiles were significantly more similar within than between YHRD metapopulations, then Y-STR profiles from the same YHRD metapopulation should ‘group’ in an MDS plot and somehow distinguish themselves from other YHRD metapopulations. However, visual inspection of the marker kit-specific MDS plots provided no evidence that this was true for the major YHRD metapopulations (Fig. 4). Only in a few cases, mainly for slowly mutating markers, was a certain clustering noticeable (e.g. for major YHRD metapopulations African or Eskimo Aleut). In all other cases, the Y-STR profiles from a given major YHRD metapopulation were approximately evenly distributed throughout the MDS plot of the entire data set. Similar results were obtained using UMAP and t-SNE analyses for dimensionality reduction (Supplementary Figs. S.3 and S.4), and when considering certain subgroups of the major YHRD metapopulations separately (Supplementary Figs. S.5-S.7).

**Fig. 4:**
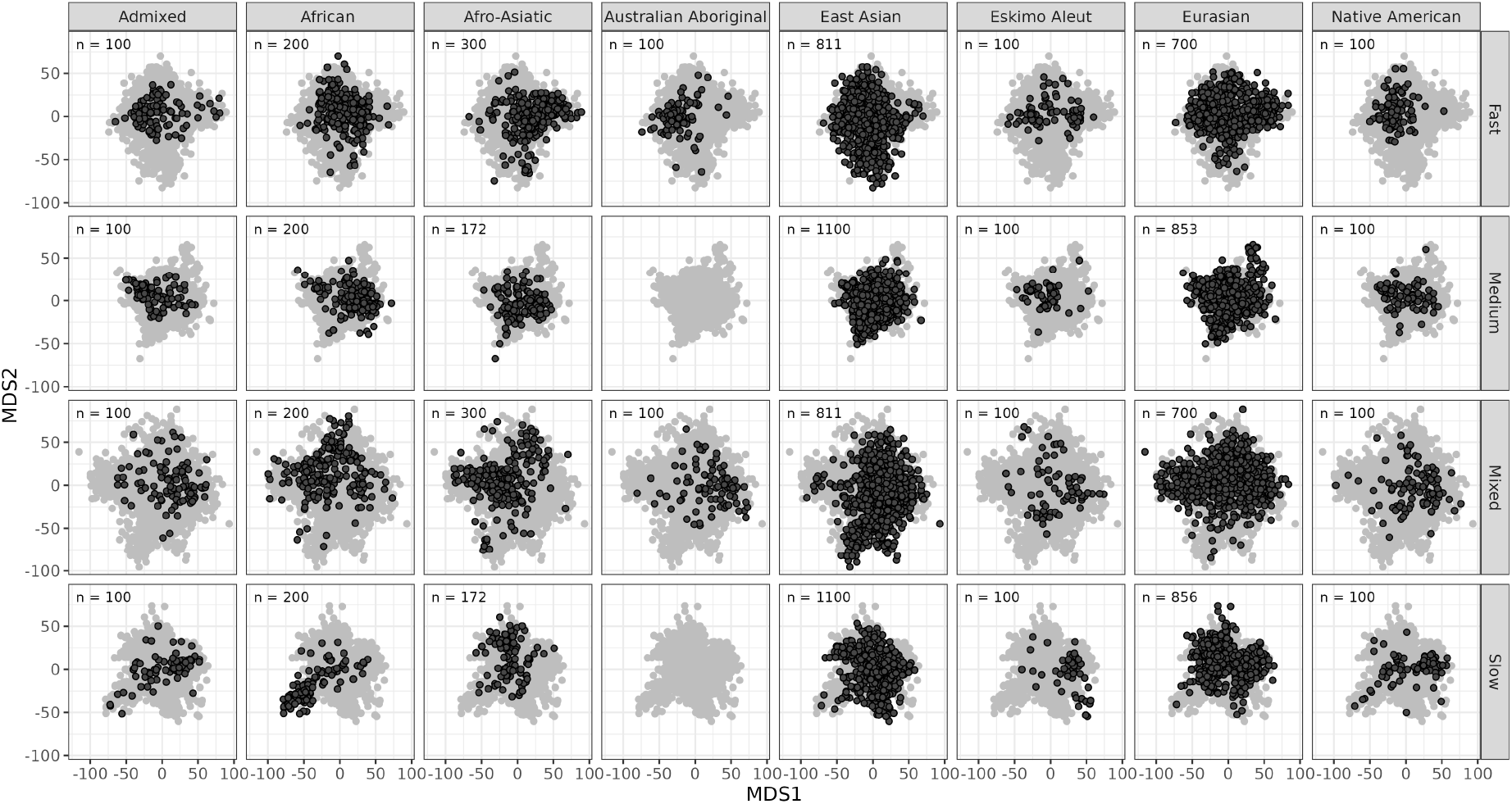
Multidimensional scaling (MDS) analysis of Y-STR profiles. In each column, the Y-STR profiles from one major YHRD metapopulation are highlighted in black. MDS1 (MDS2): first (second) MDS coordinate; *n*: number of Y-STR profiles included. For the definition of marker kits, see main text and Table 1.

### 3.4. Clustering

#### 3.4.1. Hierarchical clustering

A suitable agglomeration method for hierarchical clustering was selected for each marker kit by maximisation of the respective agglomeration coefficient *ac* (see Section 2.5). Of the five agglomeration methods considered (‘average’, ‘single’, ‘complete’, ‘Ward’, and ‘weighted’), the Ward method consistently yielded both the highest and the least variable *ac* values for the different marker kits (Table 2). Therefore, hierarchical clustering was performed applying the Ward method to distance measure *d*_1_.

For each marker kit, we determined the height of the hierarchical cluster dendrogram at which a given residual number of clusters was reduced by one by the subsequent merger. Applying the elbow method [43] to a graphical representation of these residual cluster numbers and merger dendrogram heights (Fig. 5), we assigned six, four, seven, and five structurally evident clusters to the Y-STR profiles for the fast, medium, mixed, and slow mutating marker kit, respectively. The corresponding cluster dendrograms are provided in Supplementary Fig. S.8.

**Fig. 5:**
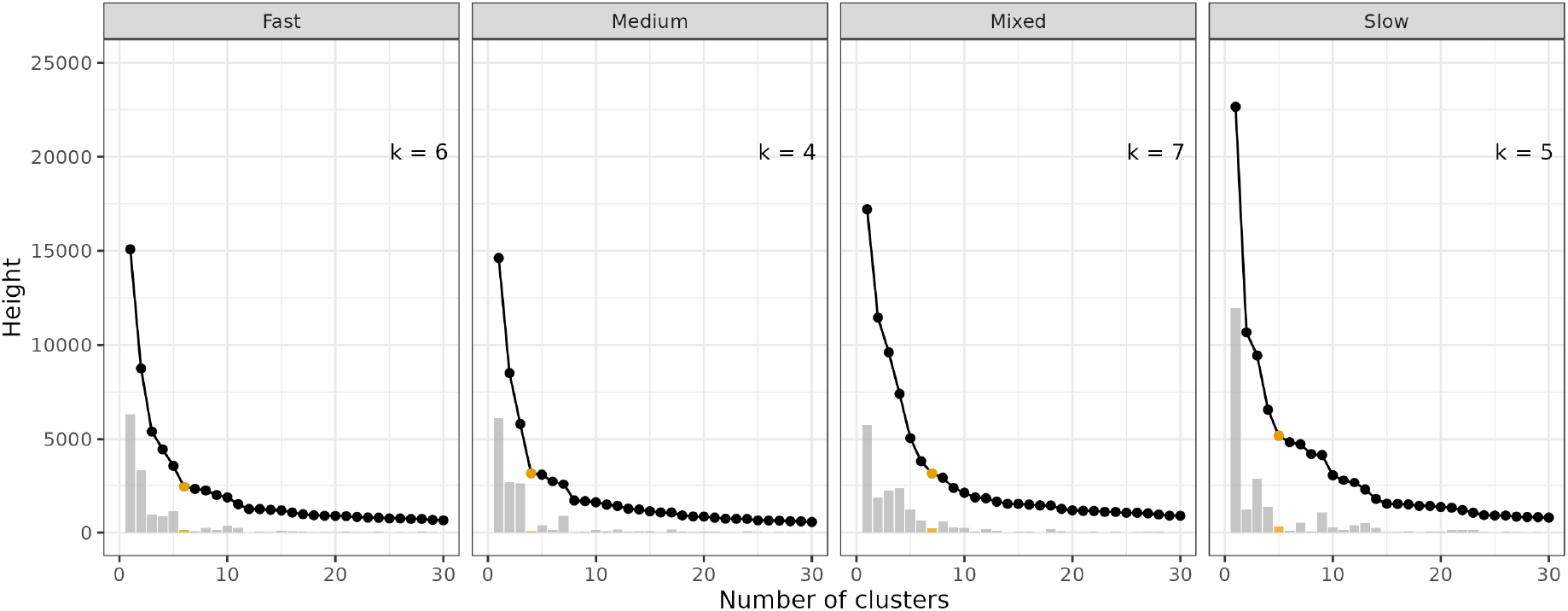
Residual cluster number and merger dendrogram height during hierarchical clustering. *k*: number of clusters present as identified by the elbow method (also highlighted in orange); grey bars: decrease in dendrogram height when the residual cluster number is increased by one. For the definition of marker kits, see main text and Table 1.

The stability of the clusters identified in this way was assessed by bootstrapping (100 bootstrap samples per marker kit), using the Jaccard index as a measure of cluster overlap. Several clusters were characterised by an average maximum Jaccard index, taken over the bootstrap samples, around or below 0.5 (Table 3) which means that, at best, half of the Y-STR profiles from the cluster were, on average, also grouped together in bootstrap clusters. Furthermore, the inferred clusters proved to be most stable for the slowly mutating marker kit which seems plausible since the allelic similarity and historical relatedness of Y-STR profiles are more likely to be correlated when mutation rates are small.

**Table 3.** Average maximum Jaccard indices obtained during bootstrapping-based assessment of cluster stability.

| Marker kit<sup>a</sup> | Structural cluster |  |  |  |  |  |  |
| --- | --- | --- | --- | --- | --- | --- | --- |
|  | 1 | 2 | 3 | 4 | 5 | 6 | 7 |
| **Fast** | 0.70 | 0.61 | 0.53 | 0.53 | 0.47 | 0.27 |  |
| **Medium** | 0.66 | 0.62 | 0.50 | 0.44 |  |  |  |
| **Mixed** | 0.65 | 0.55 | 0.50 | 0.45 | 0.43 | 0.41 | 0.29 |
| **Slow** | 0.76 | 0.75 | 0.74 | 0.64 | 0.48 |  |  |

<sup>a</sup>For the definition of marker kits, see main text and Table 1. Note: Clusters were numbered *post hoc* according to cluster stability.

We next determined how the Y-STR profiles from a given major YHRD metapopulation were distributed over the marker kit-specific clusters identified (Fig. 6). With the exception of slowly mutating markers, the clusters showed only little major YHRD metapopulation specificity or, in cases where a cluster dominated one particular metapopulation, it also did so for another, historically distant major YHRD metapopulation. These results suggest that major YHRD metapopulations would be difficult to recover by agnostic clustering of Y-STR profiles alone, at least for medium and fast mutating markers. For slowly mutating markers, in contrast, some major YHRD metapopulations overlapped significantly with specific clusters and, with the exception of Eurasians, at least 50% of each major YHRD metapopulation belonged to only a single cluster. Moreover, significant overlaps in cluster membership as, for example, between Eskimo Aleut and Native Americans usually appeared to be historically meaningful.

**Fig. 6:**
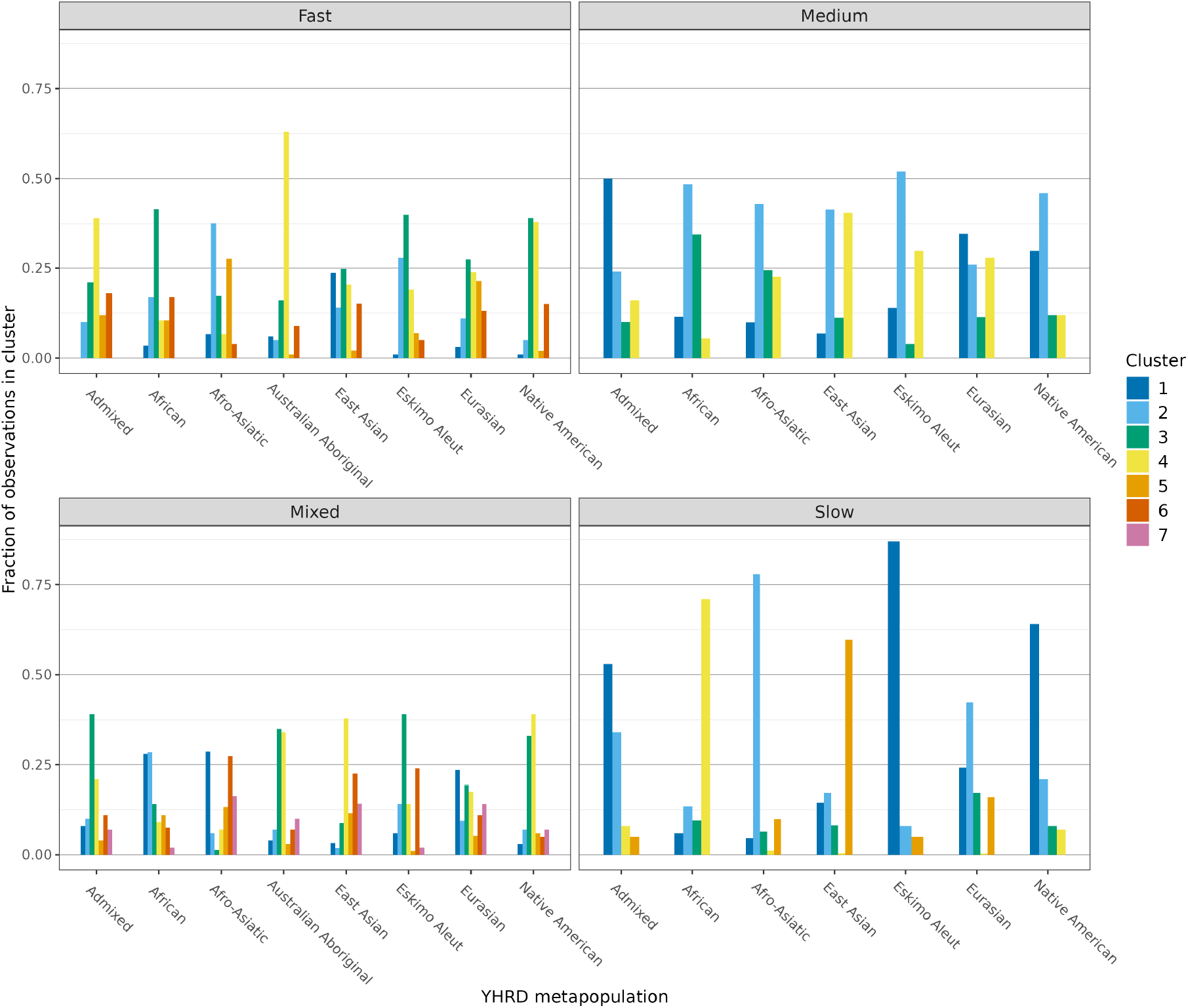
Distribution of major YHRD metapopulations over identified Y-STR profile clusters. The height of each bar corresponds to the fraction of Y-STR profiles in a given major YHRD metapopulation that belongs to a certain cluster. For the definition of marker kits, see main text and Table 1.

#### 3.4.2. STRUCTURE analysis

Y-STR profiles were also grouped by STRUCTURE [47] analysis as described in [48, 49]. To determine the number of Y-STR profile groups present, we applied the elbow method [43] to a cross-entropy criterion [49] each time calculated for a fixed group number of between one and 15. Seven, three, nine, and ten groups were identified this way for the fast, medium, mixed, and slow marker kit, respectively. Each Y-STR profile was then assigned to the group with the highest probability of membership for that profile. Line plots of the STRUCTURE results (Supplementary Fig. S.9) highlighted that many Y-STR profiles were characterised by a large probability of membership in one group. In particular, ca. 79%, 77%, 69%, and 69% of YHRD profiles belonged to one group with probability ≥ 0.9 for the fast, medium, mixed, and slow marker kit, respectively.

Similar to hierarchical clustering, STRUCTURE yielded groupings of Y-STR profiles that, in most cases, only poorly matched the definition of major YHRD metapopulations (Fig. 7). Only a few major YHRD metapopulations had more than 50% of their Y-STR profiles assigned to one and the same group. In view of the large number of groups inferred for the mixed and slow marker kits, we extended the analysis for these marker kits so as to also cover the predefined subgroups of major YHRD metapopulations (Supplementary Fig. S.10). Since STRUCTURE assigns Y-STR profiles to groups only with a certain probability, we additionally repeated the STRUCTURE analyses only for profiles that had a group membership probability ≥ 0.9 (Supplementary Fig. S.11). Both analyses yielded similar results to the original.

**Fig. 7:**
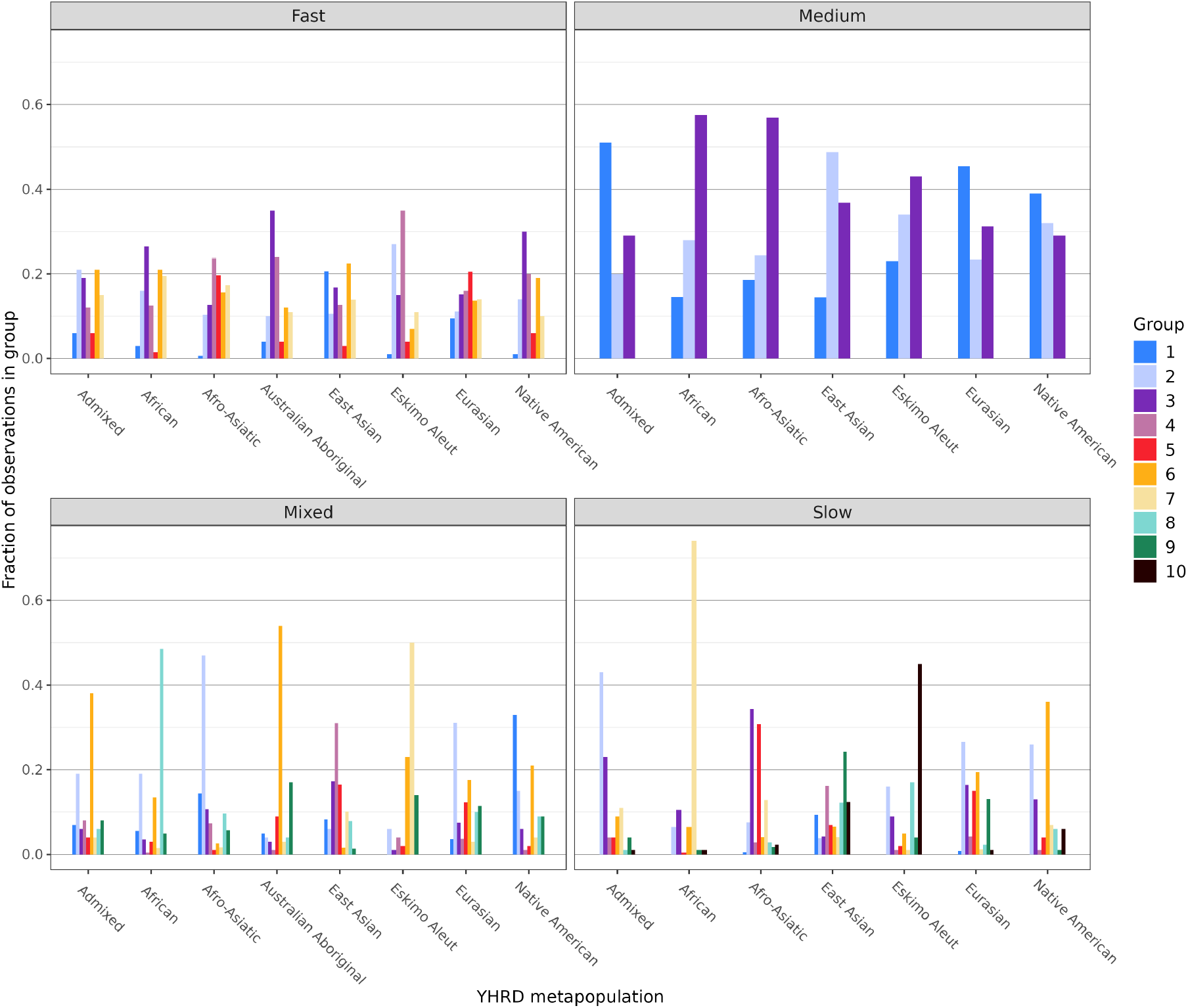
Grouping of Y-STR profiles by STRUCTURE analysis. The height of each bar corresponds to the fraction of profiles in a major YHRD metapopulation that belongs to a certain STRUCTURE-derived group. For the definition of marker kits, see main text and Table 1.

## 4. Discussion

When equating the pairwise distance between Y-STR profiles to the sum of marker-wise allele differences, classical cluster analysis suggests only weak inherent structure in the data currently included in YHRD. Furthermore, some of the clusters identified in our study proved unstable when judged by the Jaccard indices obtained from comprehensive bootstrapping. Notably, clusters derived for Y-STR profiles comprising slowly mutating markers exhibited greater stability than others, a pattern consistent with previous observations for much smaller numbers of markers [50–52]. Also consistent with other studies [37], the most pronounced differences in YHRD were observed between Y-STR profiles of African and non-African origin, while Eurasian profiles tended to be most similar to one another.

Our results suggest that YHRD cannot be divided into clearly distinguishable subsamples that correspond strongly to the metapopulations highlighted by YHRD itself, at least not by drawing solely upon the Y-STR profiles themselves. On the contrary, the relationship between Y-STR haplotype and metapopulation membership proved to be highly variable and dependent upon the markers considered. The metapopulations highlighted in YHRD are therefore not well-defined genetic units. This finding does not call into question the concept of metapopulations per se, but rather suggests that the human genetic clock of the Y-chromosome has ticked too differently from cultural and geographical clocks for the traces of these clocks to run synchronously.

What does this mean for forensic practice, support of which is one of the essential if not the most essential concerns of YHRD? Some of us have previously pointed out that equating database frequency estimates with match probabilities in forensic casework requires that the database, or at least the part of it that is being used, represents a meaningful case-specific suspect population [5]. At first glance, it seems obvious that the definition of this population should not be based primarily upon genetic considerations to avoid circular reasoning. Instead, it is generally accepted that the group of alternative trace donors is first narrowed down using non-genetic information (location, time, cultural context, witness statements, etc.) and only then characterized in detail genetically [2]. In this second step, it can of course be useful to favor genetic markers that distinguish the suspect population from other groups of males.

These considerations inevitably lead to the question of whether and under what circumstances a YHRD metapopulation can be considered a plausible suspect population. First of all, “plausible” in this context must mean that the definition of the metapopulation applies to the suspect population in question almost unchanged. Furthermore, this applicability must be fully accepted by all parties involved in the case, including the defense. In our view, the whole approach is therefore only viable if the criteria used to define YHRD metapopulations are made fully transparent in advance to all stakeholders which, to our knowledge, has not yet been the case.

In view of the general problems associated with the forensic use of population database information [5], alternative approaches to evaluate trace-donor matches of Y-STR profiles have been proposed, including evolutionary simulation of potential matches [4] and calculation of the exact match probability inside and outside the suspect pedigree [53]. However, it will certainly take some time before such novel approaches can be incorporated into forensic casework, and the current practice of estimating profile frequencies using databases is therefore likely to be maintained for a while. If this is the case, then the genetic information contained in YHRD must at least be used to select metapopulations - to serve as suspect populations - in a way that is fair to the suspects while preserving the evidentiary value of the database as much as possible. A two-step approach is conceivable for this.

First, the YHRD metapopulation with the (likely) strongest Y-chromosomal similarity to the suspect is identified after restricting the Y-STR profile to the slowly mutating markers. Developing a suitable method for such a selection admittedly requires further research. Be that as it may, if the selection is consistent with the non-genetic evidence for a particular suspect population, it can also be considered fair to the suspect. “Fair” here refers to the principle of “in dubio pro reo” (presumption of innocence) because a suspect profile generated for any other set of Y-STR markers should also be quite frequent, if not most frequent, in the selected metapopulation. In the second step, the frequency of the (much more suspect-specific) Y-STR profile comprising moderately to rapidly mutating markers is estimated in the selected YHRD metapopulation and, if this is to be practiced despite theoretical reservations, equated to the match probability. In the vast majority of cases, the persuasiveness of the result of the second step should be similar to that of using the entire profile. Furthermore, since mutations at different Y-STRs are statistically independent, the use of different markers for the initial identification and subsequent genetic characterization of a suspect population would also help to avoid circular reasoning when evaluating trace-donor profile matches. Should the genetic and non-genetic information point to very different suspect populations in step one, we recommend applying the approach to both selections and presenting both results in court.

Although our fundamental reservations about the approach taken so far for obtaining match probabilities remain, and although YHRD offers only limited coverage for many populations, particularly Non-Eurasians [54], we believe that a change to the procedure as proposed here could represent a significant improvement in terms of its scientific and legal irrefutability.

## Appendix A. Distance properties of distance measures *d*_1_, *d*_2_, and *d*_3_

Any valid distance measure, *d*, must fulfil the following properties for Y-STR profiles *x*_1_, *x*_2_, and *x*_3_:

1. *d*(*x*_1_, *x*_2_) = 0 ⇔ *x*_1_ = *x*_2_ (Reflexivity).
2. *d*(*x*_1_, *x*_2_) ≥ 0 (Non-negativity).
3. *d*(*x*_1_, *x*_2_) = *d*(*x*_2_, *x*_1_) (Symmetry).
4. *d*(*x*_1_, *x*_2_) + *d*(*x*_1_, *x*_3_) ≥ *d*(*x*_2_, *x*_3_) (Triangle inequality).

### Measure *d*_1_

For *d*_1_, the first three properties follow from the fact that (i) log(*µ*_*j*_) *<* 0 since *µ*_*j*_ ∈ (0, 1), (ii) *m*_*j*_ ≥ 0 and *m*_*j*_ = 0 for all *j* if and only if the two Y-STR profiles in question are identical, and (iii) *m*_*j*_ is itself symmetric.

Let 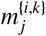 denote the absolute allelic difference between *x*_*i*_ and *x*_*k*_ at the *j*^th^ locus. Then

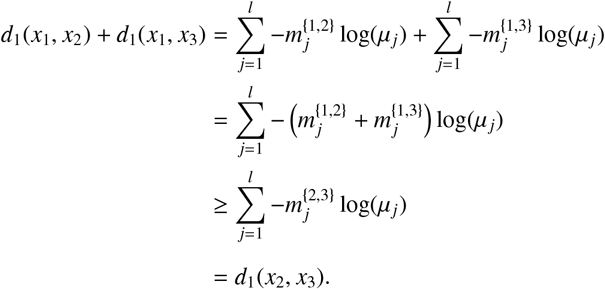

Thus, the triangle inequality also holds for *d*_1_.

### Measure *d*_2_

Measure *d*_2_ satisfies the first three properties of a distance measure for the same reasons as *d*_1_ (see above). The triangle inequality holds for *d*_2_ because

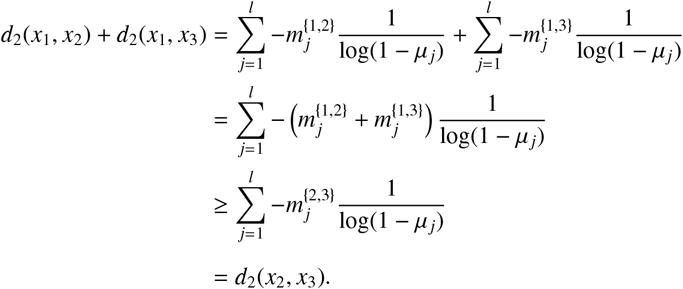

### Measure *d*_3_

Reflexivity and symmetry hold for *d*_3_ by definition. Further, even for rapidly mutating Y-STRs, it appears reasonable to assume that *N*_*e*_ ≥ 1*/*(1 − 2*µ*_*j*_). Hence, 2*µ*_*j*_ + 1*/N*_*e*_ ≤ 1 and *m*_*j*_ + 1 ≥ 1, which implies non-negativity. Finally, the triangle inequality also holds for *d*_3_ because if 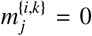 for any *j, i*, and *k*, then 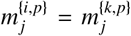 for any *p*, and if 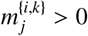 for all *j, i*, and *k*, then

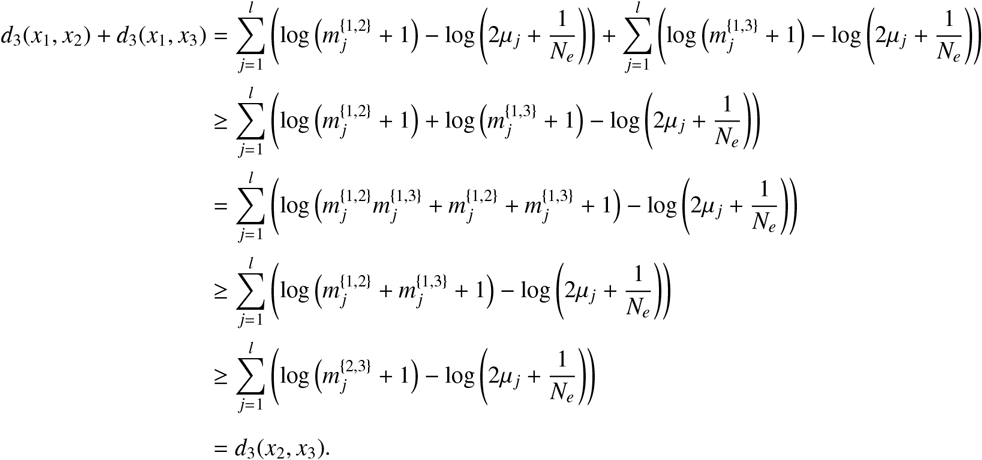

## Appendix B. Motivation of distance measure *d*_3_

The following considerations are based upon [55, 56]. Let *x*_1_ and *x*_2_ be two single-locus Y-STR profiles for which their coalescence time *T* should be estimated by some meaningful value 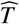 . Let *m*_*i*_, for *i* ∈ {1, 2}, denote the number of mutations that *x*_*i*_ has undergone since the most recent common ancestor of *x*_1_ and *x*_2_.

As an estimate 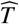, we want to derive the posterior expectation of *T* given the allele difference between *x*_1_ and *x*_2_. To this end, we assume the following prior distribution for *T* :

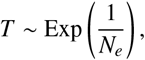

where *N*_*e*_ is the effective population size [56]. If *µ*denotes the mutation rate, then the conditional distribution of *m*_*i*_ given *T* equals

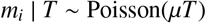

for *i* ∈ {1, 2}. Hence, the posterior distribution of *T* is given by

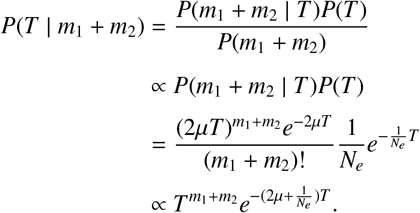

This means that 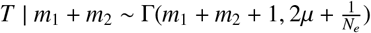 with (posterior) expectation

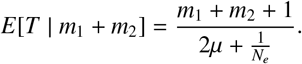

In case of *l* loci, the likelihood equals the product of the locus-specific terms so that the posterior expectation now equals

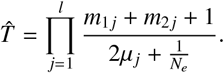

To avoid numerical overflow, we will take

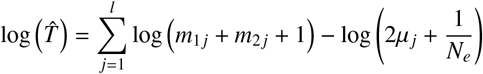

and hence, we propose distance measure

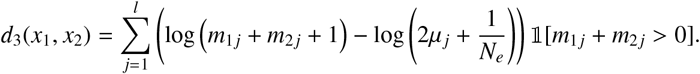

Pragmatically, we set *m*_1 *j*_ + *m*_2 *j*_ equal to the absolute allelic difference between *x*_1_ and *x*_2_ at the *j*^th^ locus, thereby knowingly disregarding the possibility of (undetectable) backwards mutations.

