## Supplementary material for "Are genetically defined “metapopulations” self-evident in YHRD?"

**Table S.1:** Number of Y-STR profiles per YHRD metapopulation and marker kit.

| YHRD metapopulation | Fast | Medium | Mixed | Slow |
|---|---|---|---|---|
| Admixed | 6,563 | 7,712 | 6,750 | 7,668 |
| African - African American | 2,624 | 4,494 | 2,587 | 4,432 |
| African - Sub-Saharan African | 2,096 | 2,313 | 2,056 | 2,291 |
| Afro-Asiatic - Berber | 281 | 0 | 291 | 0 |
| Afro-Asiatic - Cushitic | 379 | 72 | 394 | 72 |
| Afro-Asiatic - Semitic | 1,420 | 2,683 | 2,158 | 2,676 |
| Australian Aboriginal | 732 | 0 | 740 | 0 |
| East Asian | 2,955 | 2,958 | 2,988 | 2,934 |
| East Asian - Austro-Asiatic | 2,107 | 2,794 | 2,105 | 2,792 |
| East Asian - Austronesian | 198 | 1,640 | 198 | 1,618 |
| East Asian - Dravidian | 11 | 245 | 11 | 321 |
| East Asian - Japanese | 1,287 | 322 | 1,283 | 322 |
| East Asian - Korean | 1,296 | 2,430 | 1,314 | 2,937 |
| East Asian - Sino-Tibetan - Chinese (Han) | 46,705 | 26,613 | 46,635 | 31,338 |
| East Asian - Sino-Tibetan - Tibeto-Burman | 7,338 | 6,677 | 7,329 | 6,646 |
| East Asian - Tai-Kadai | 5,775 | 3,310 | 5,779 | 3,298 |
| Eskimo Aleut | 189 | 288 | 189 | 285 |
| Eurasian - Altaic | 4,920 | 2,443 | 4,865 | 2,338 |
| Eurasian - European | 2,872 | 4,780 | 2,928 | 4,769 |
| Eurasian - European - Eastern European | 2,129 | 1,883 | 2,150 | 1,873 |
| Eurasian - European - South-Eastern European | 1,013 | 3,504 | 1,027 | 3,448 |
| Eurasian - European - Western European | 4,541 | 15,461 | 4,580 | 15,399 |
| Eurasian - Indian | 1,216 | 4,289 | 1,219 | 4,369 |
| Eurasian - Indo-Iranian | 714 | 956 | 660 | 953 |
| Native American | 2,892 | 3,370 | 2,893 | 3,323 |
| East Asian - Indo-Pacific | 0 | 162 | 0 | 163 |
| East Asian - Sino-Tibetan | 0 | 142 | 0 | 140 |
| Eurasian - Caucasian | 0 | 53 | 0 | 56 |
| Eurasian - Uralic-Yukaghir | 0 | 1,286 | 0 | 1,287 |

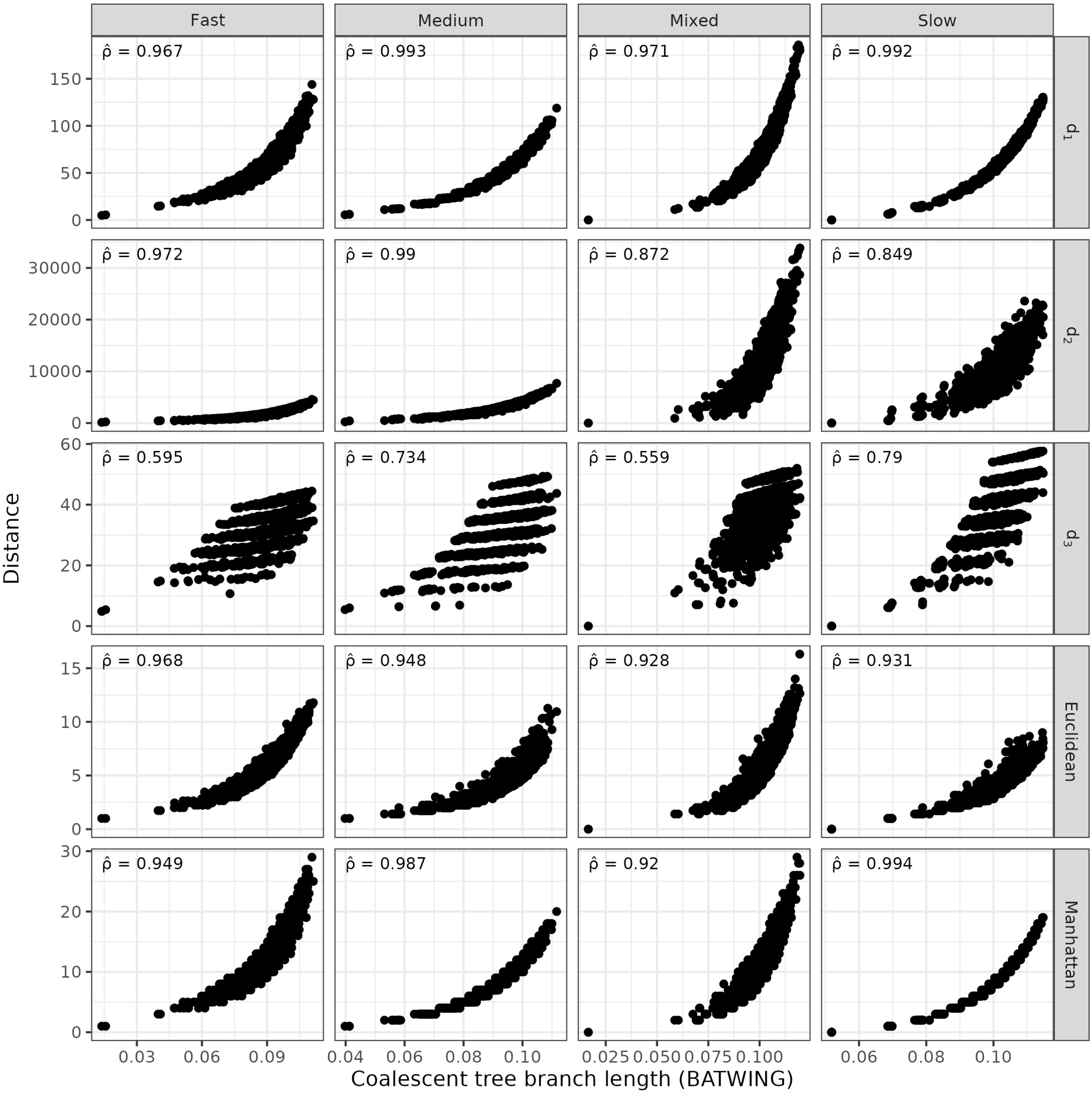


**Fig. S.1:** Benchmarking of pairwise Y-STR profile distance measures against BATWING. For each pair of Y-STR profiles, the respective distance is plotted against the coalescent branch length estimated with BATWING, assuming an effective population size of 5,000. $\hat{\rho}$: average Spearman's rank correlation coefficient, taken over five repetitions of the analysis. For the definition of marker kits, see main text and Table 1.

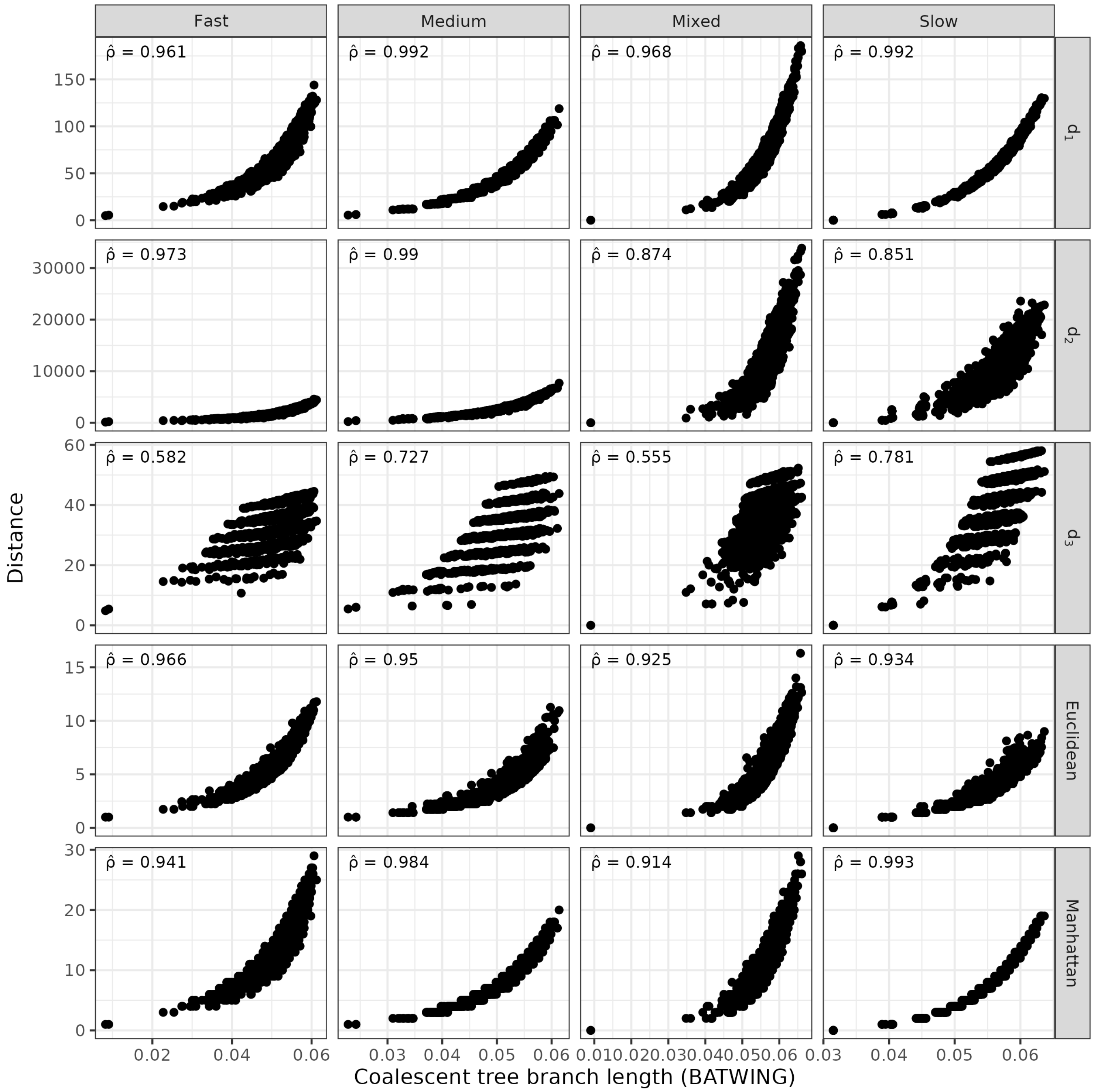


**Fig. S.2:** Benchmarking of pairwise Y-STR profile distance measures against BATWING. For each pair of Y-STR profiles, the respective distance is plotted against the coalescent branch length estimated with BATWING, assuming an effective population size of 10,000. $\hat{\rho}$: average Spearman's rank correlation coefficient, taken over five repetitions of the analysis. For the definition of marker kits, see main text and Table 1.

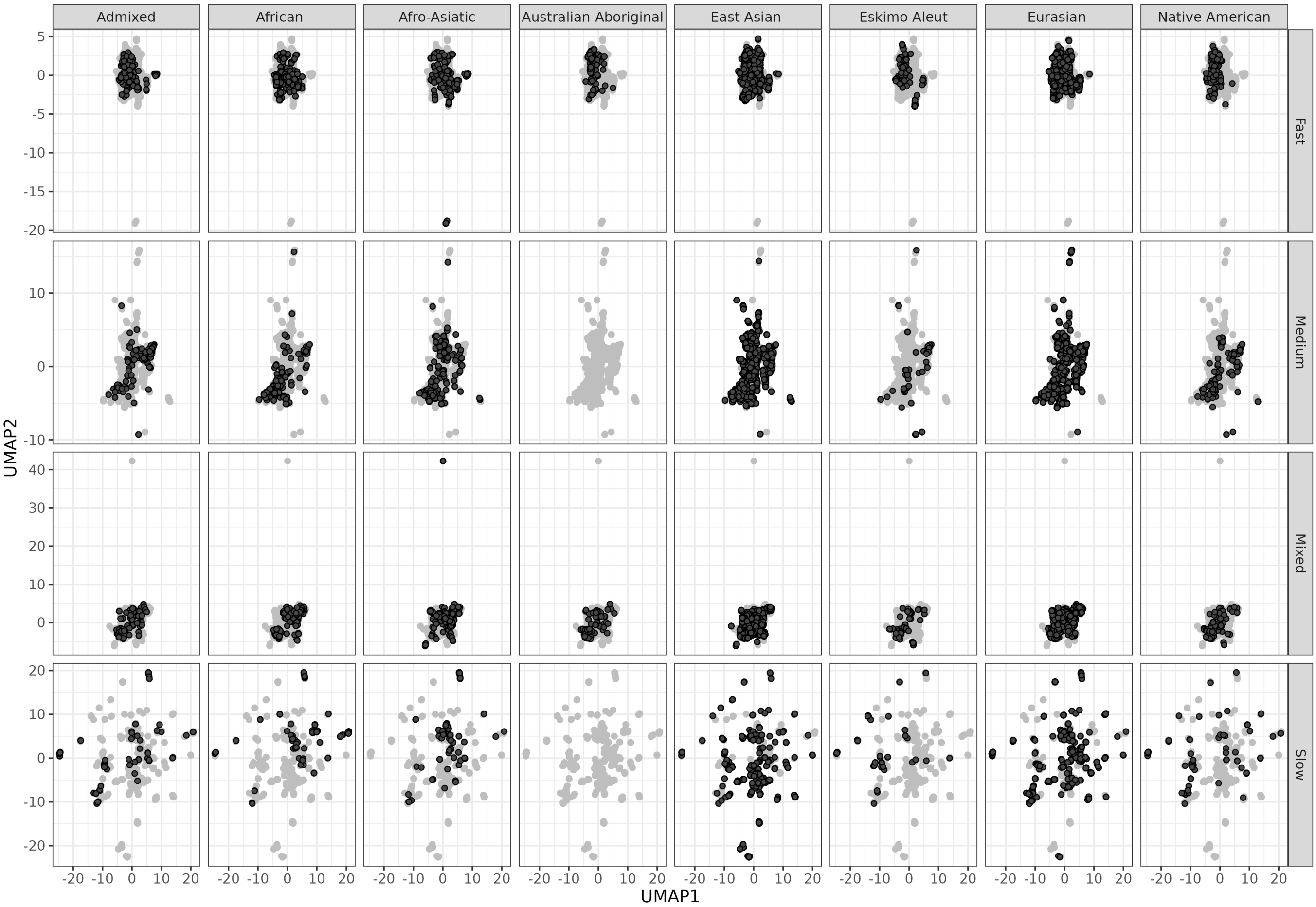


**Fig. S.3:** Uniform Manifold Approximation and Projection (UMAP) analysis of Y-STR profiles. In each column, the Y-STR profiles from one major YHRD metapopulation are highlighted in black. UMAP1 (UMAP2): first (second) UMAP coordinate. For the definition of marker kits, see main text and Table 1.

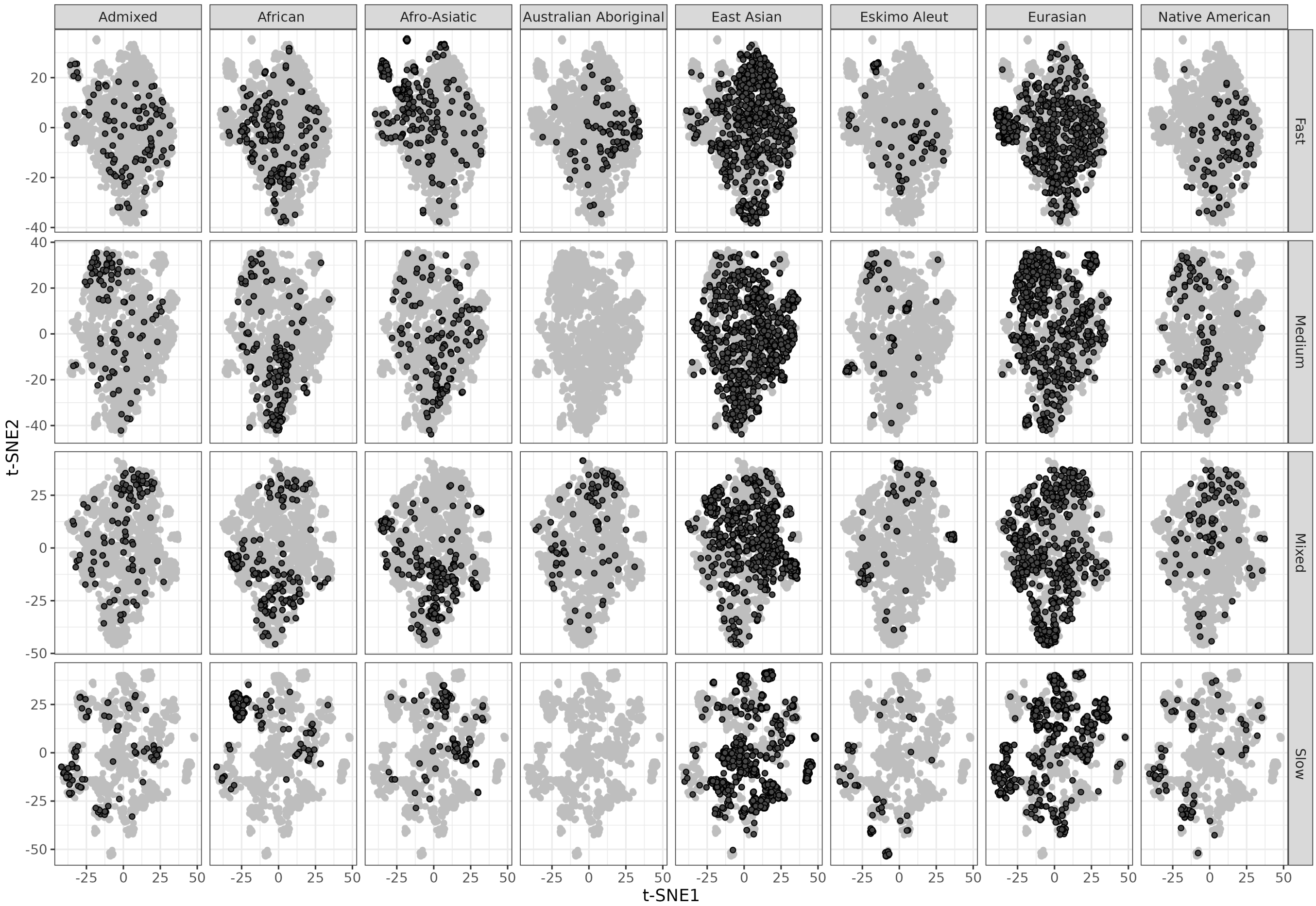


**Fig. S.4:** t-Distributed Stochastic Neighbor Embedding (t-SNE) analysis of Y-STR profiles. In each column, the Y-STR profiles from one major YHRD metapopulation are highlighted in black. t-SNE1 (t-SNE2): first (second) t-SNE coordinate. For the definition of marker kits, see main text and Table 1.

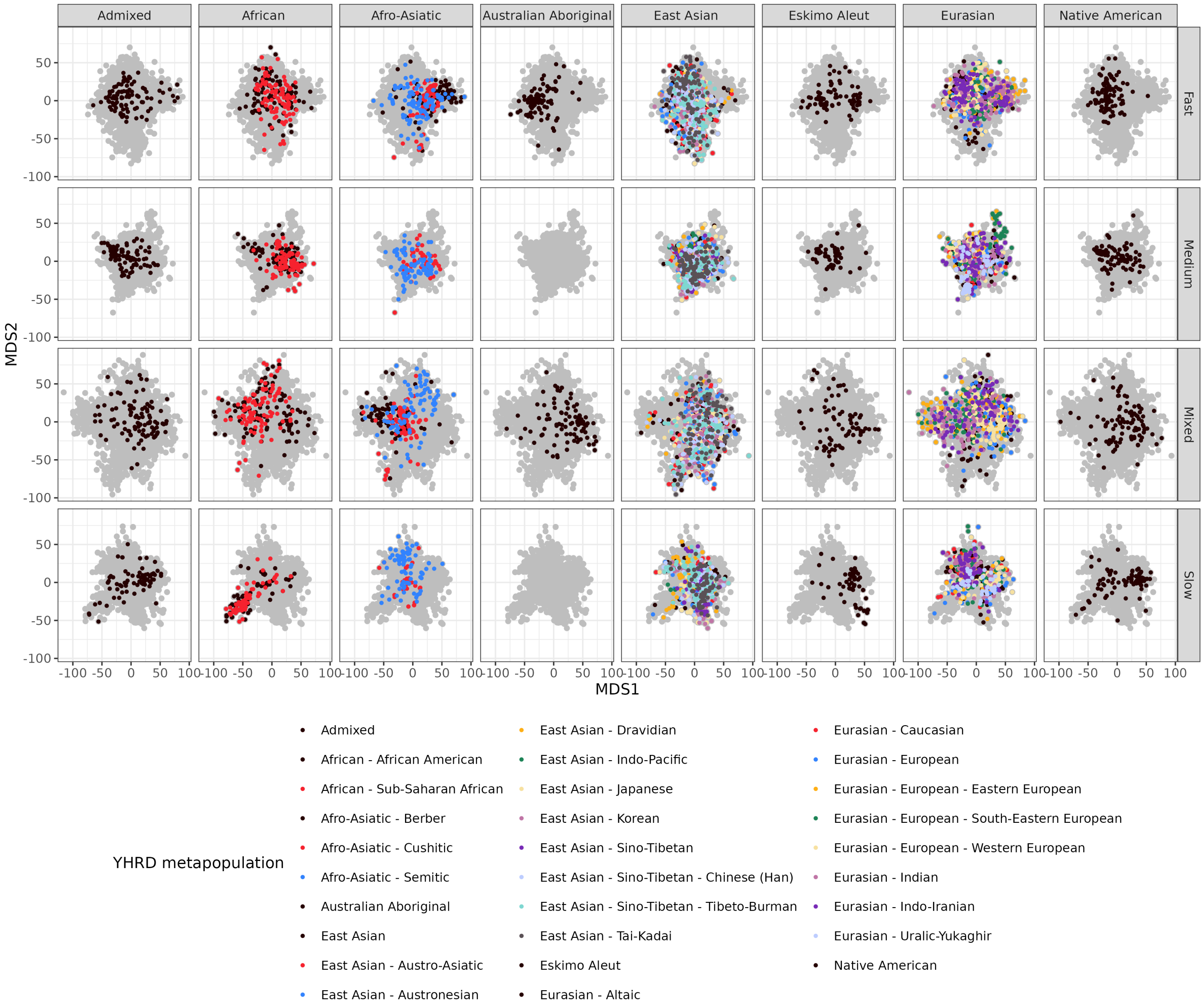


**Fig. S.5:** MDS analysis of Y-STR profiles. In each column, the Y-STR profiles from subgroups of one major YHRD metapopulation are highlighted. MDS1 (MDS2): first (second) MDS coordinate. For the definition of marker kits, see main text and Table 1.

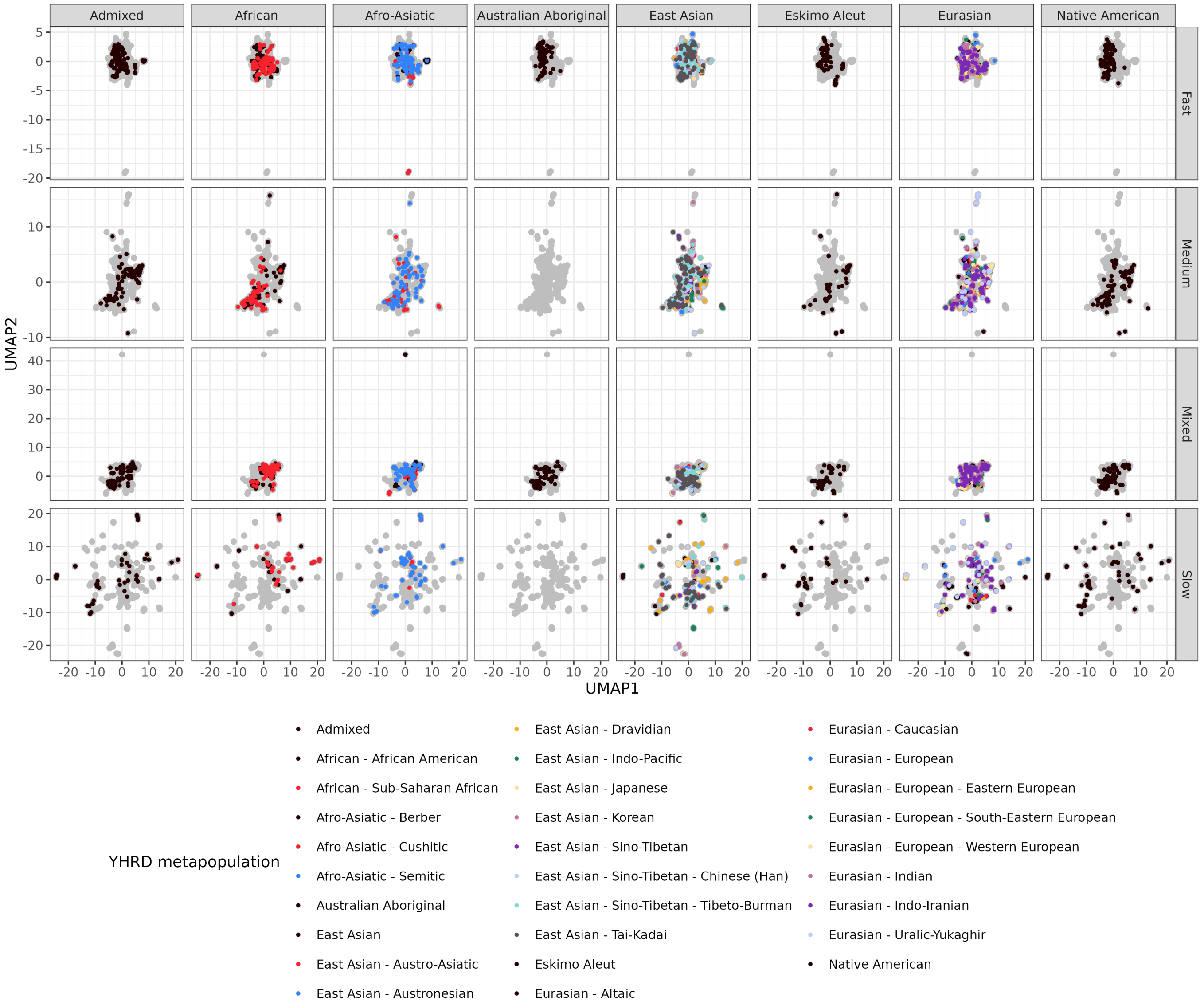


**Fig. S.6:** UMAP analysis of Y-STR profiles. In each column, the Y-STR profiles from subgroups of one major YHRD metapopulation are highlighted. UMAP1 (UMAP2): first (second) UMAP coordinate. For the definition of marker kits, see main text and Table 1.

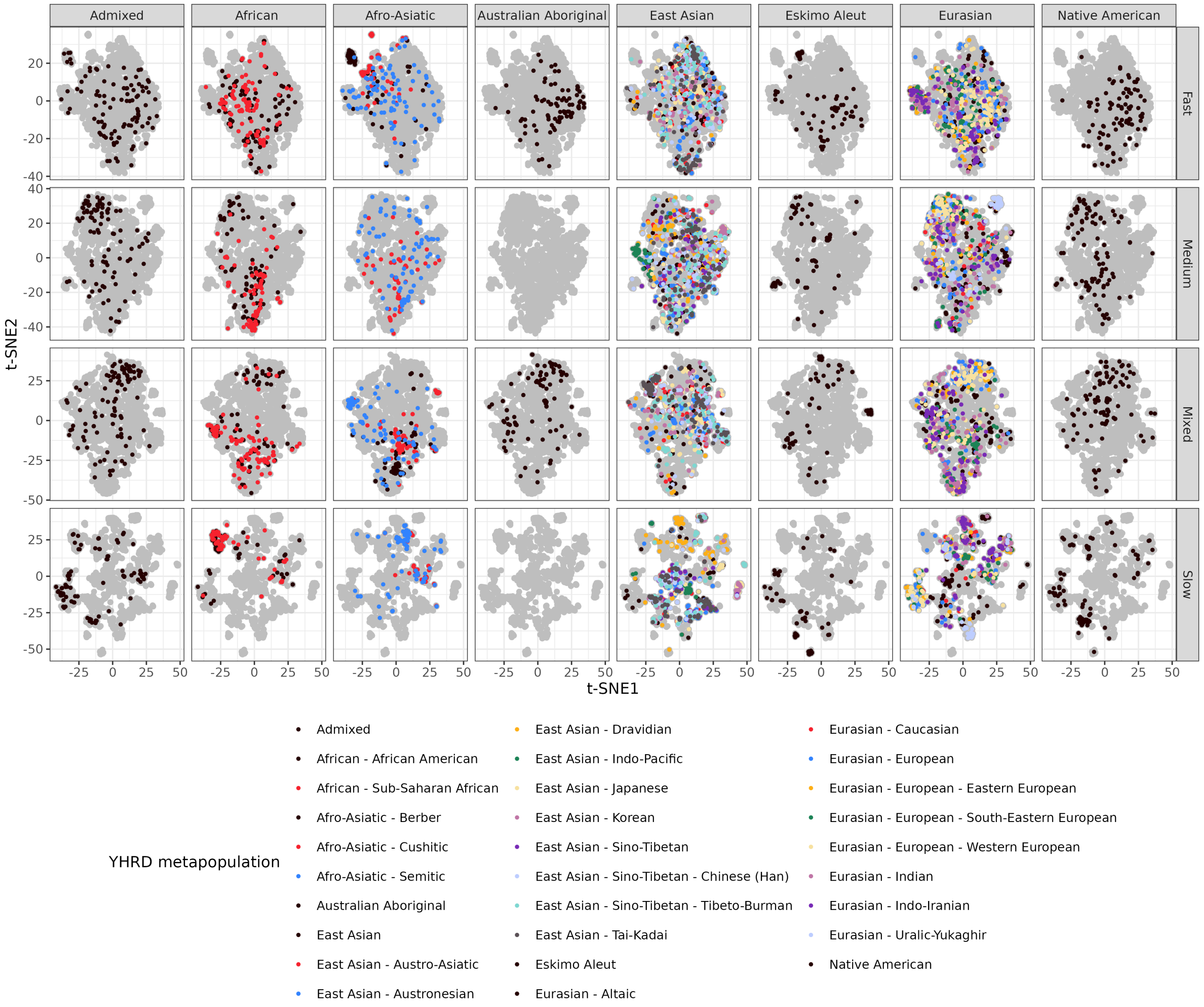


**Fig. S.7:** t-SNE analysis of Y-STR profiles. In each column, the Y-STR profiles from subgroups of one major YHRD metapopulation are highlighted. t-SNE1 (t-SNE2): first (second) t-SNE coordinate. For the definition of marker kits, see main text and Table 1.

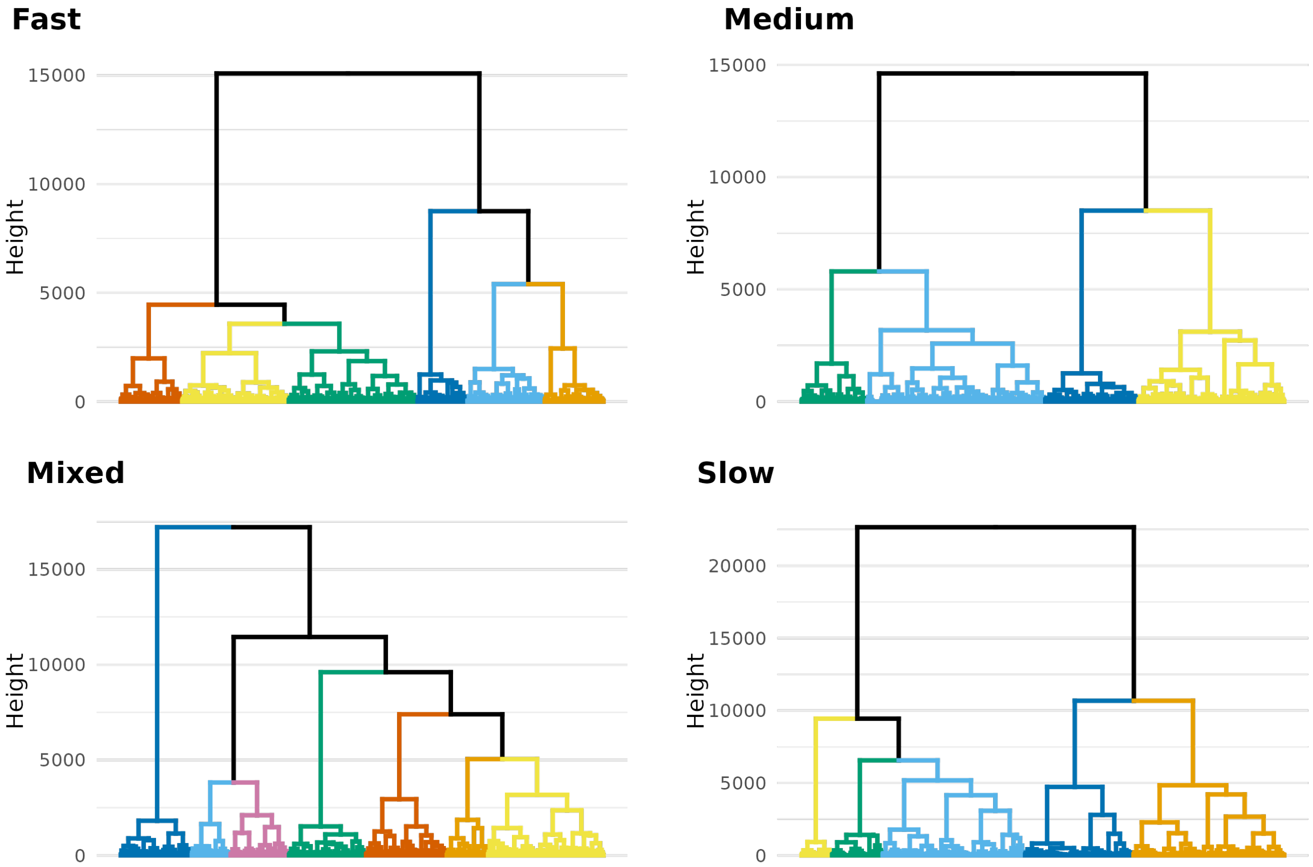


**Fig. S.8:** Dendrograms for the hierarchical clustering analysis for each marker kit. The clusters from the hierarchical clustering analysis are highlighted according to the same coloring scheme as in Fig. 6.

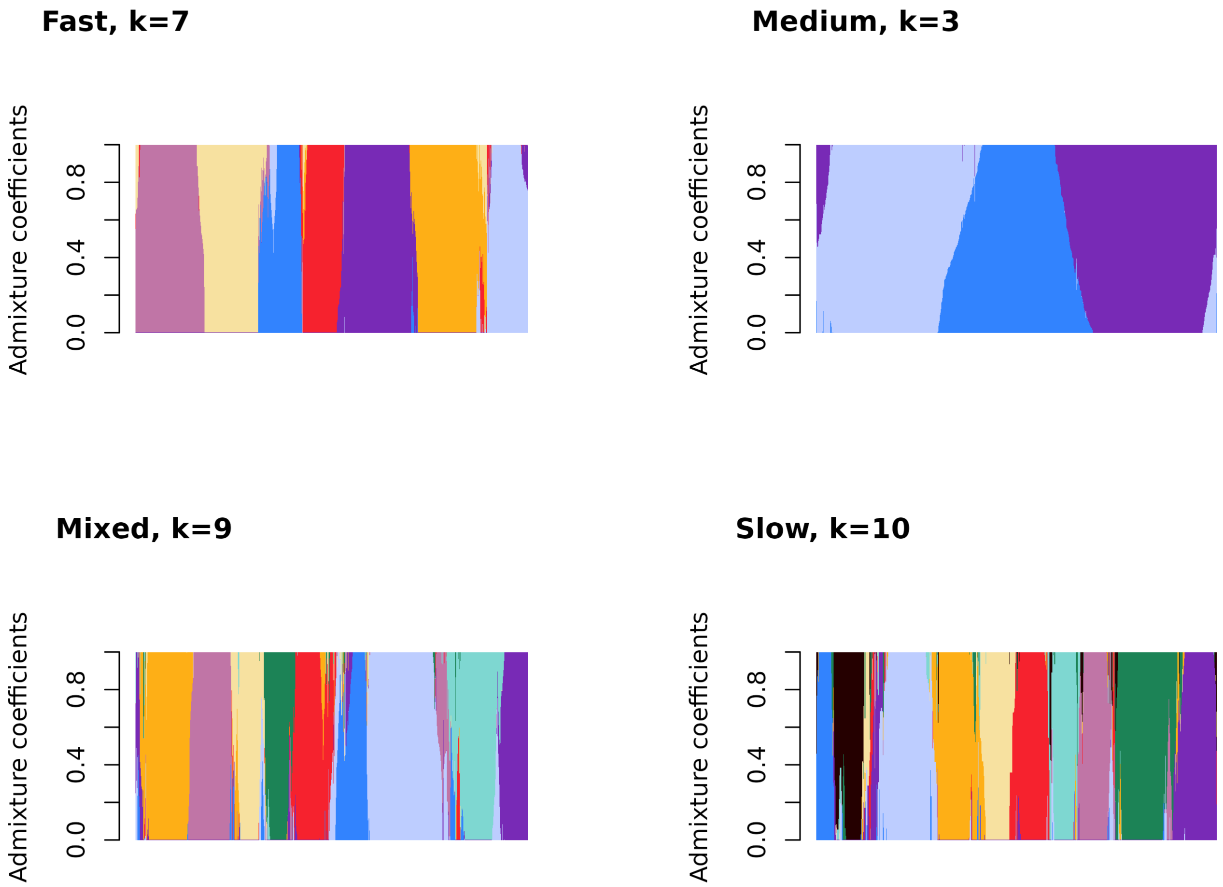


**Fig. S.9:** Line plots of STRUCTURE-inferred admixture in YHRD for each marker kit. Each vertical line corresponds to a YHRD profile. The line colouring illustrates the admixture coefficients that measure, for each profile, the relative likelihood of belonging to one of the $k$ groups inferred for the respective marker set. The $k$ groups are highlighted using the same coloring scheme as in Fig. 7.

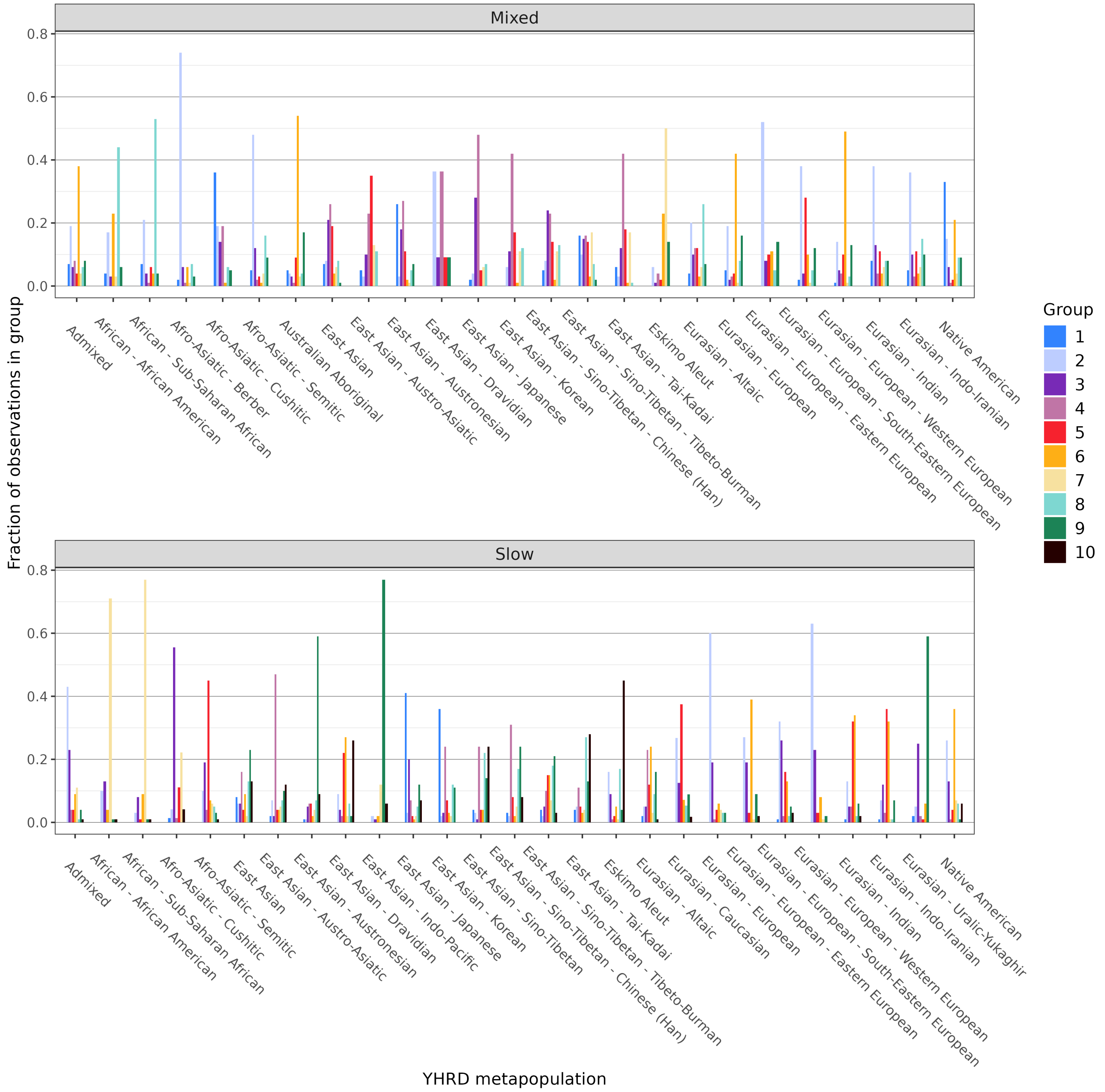


**Fig. S.10:** Grouping of Y-STR profiles by STRUCTURE analysis. The height of each bar corresponds to the fraction of profiles in a given YHRD metapopulation that belongs to a certain STRUCTURE-derived group. For the definition of marker kits, see main text and Table 1.

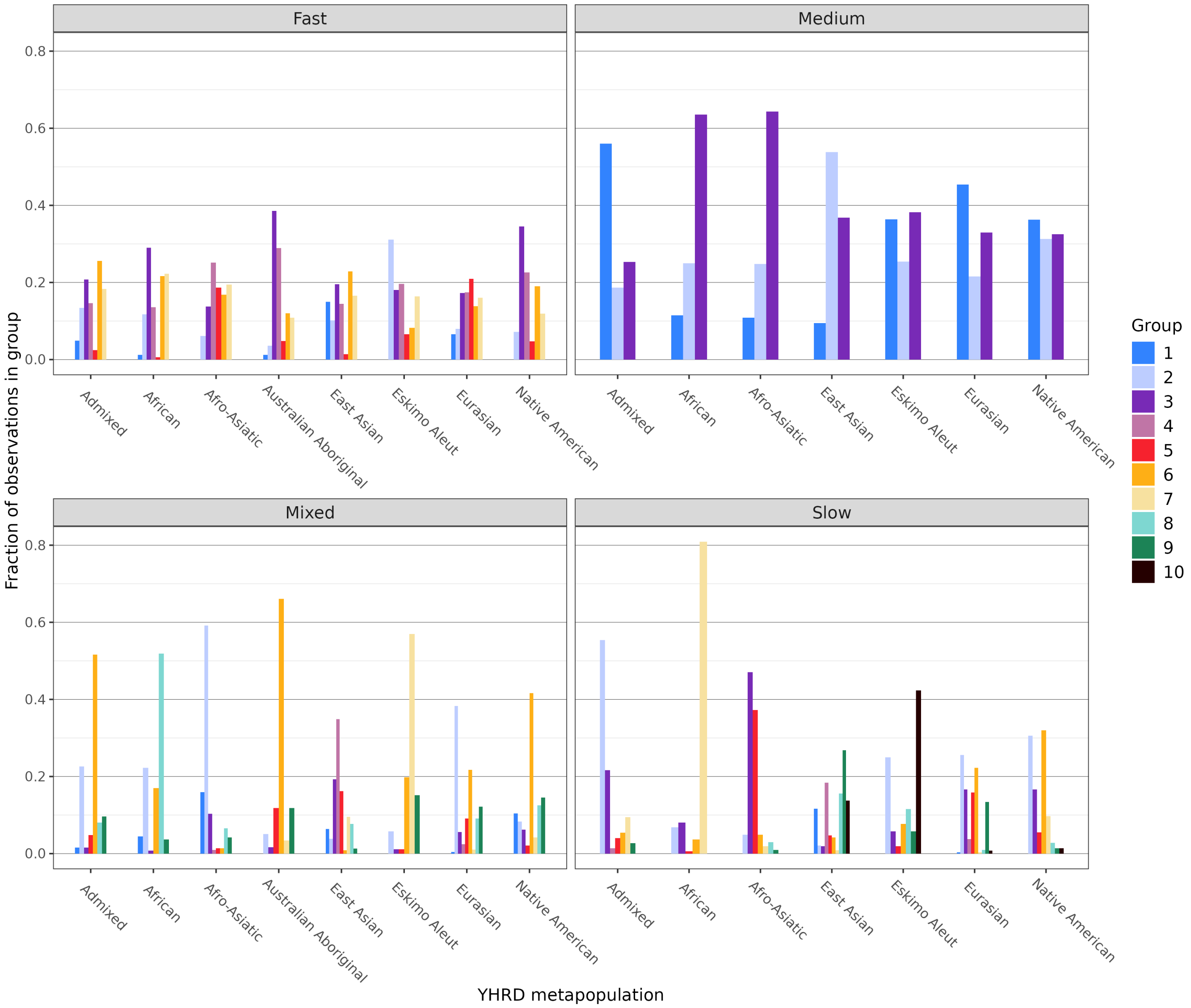


**Fig. S.11:** Grouping of Y-STR profiles by STRUCTURE analysis. The height of each bar corresponds to the fraction of profiles in a given major YHRD metapopulation that belongs to a certain STRUCTURE-derived group when only considering profiles with a probability of at least 0.9 of belonging to the assigned STRUCTURE-derived group. For the definition of marker kits, see main text and Table 1.
